# LimbLab: Pipeline for 3D Analysis and Visualisation of Limb Bud Gene Expression

**DOI:** 10.1101/2025.04.08.647723

**Authors:** Laura Aviñó-Esteban, Heura Cardona-Blaya, Marco Musy, Antoni Matyjaszkiewicz, James Sharpe, Giovanni Dalmasso

## Abstract

**Motivation:** Although some aspects of limb development can be treated as a 2D problem, a true understanding of the morphogenesis and patterning requires 3D analysis. Since the data on gene expression patterns are largely static 3D image stacks, a major challenge is an efficeint pipeline for staging each data-set, and then aligning and warping the data into a standard atlas for convenient visualisation.

**Results:** We present a novel bioinformatic pipeline tailored for 3D visualization and analysis of developing limb buds. The pipeline integrates key steps such as data acquisition, volume cleaning, surface extraction, staging, alignment, and advanced visualization techniques. Its modular design allows researchers to customize workflows while maintaining compatibility with tools such as Fiji and Vedo. Our results highlight the pipeline’s effectiveness in managing complex 3D gene expression data, enhancing the accuracy and reproducibility of limb development studies.

**Availability and Implementation:** LimbLab is released under the MIT license. The pipeline is implemented in Python and is available as an open-source package. It can be easily installed via pip and is accompanied by comprehensive documentation (https://limblab.embl.es/) to support users at various levels of expertise. The pipeline can be accessed and downloaded at https://github.com/LauAvinyo/limblab.

## Introduction

The spatial organization of cells and their interactions is crucial to elucidate various biological processes, including development, differentiation, and disease progression [14]. Understanding the development of tissues and organs often requires analysis of complex three-dimensional (3D) structures, as two-dimensional (2D) data do not capture key architectural and geometric features [24, 31]. For example, gene expression patterns, when captured in 2D, may fail to accurately represent the spatial relationships between genes and other molecular entities [6]. This issue is particularly evident in the development of the mouse limb, where spatial context is crucial [27]. Although the limb can sometimes be simplified to a 2D problem, such reductions can create false impressions of gene expression patterns and their interactions.

Advances in imaging technologies and computational methods have facilitated the collection of large amounts of 3D data. High-resolution imaging techniques, such as confocal microscopy [10], Optical Projection Tomography (OPT) [22], and Magnetic Resonance Imaging (MRI) [9], allow researchers to acquire detailed 3D datasets of tissues and organs [13, 22, 29, 30]. These technologies generate comprehensive spatial maps of gene expression, revealing the intricate details of biological structures in their native 3D context. This wealth of data provides an invaluable resource for accurately mapping gene expression and understanding the dynamics of developing limb buds and other tissues.

In species such as zebrafish and Drosophila, which develop externally, it is possible to use time-lapse imaging to effectively make movies of the entire developmental process *in vivo* and in real-time [28, 20]. In contrast, this approach remains challenging for embryos that develop internally, particularly after the gastrulation stage [5]. *In vitro* culture systems face significant limitations in recapitulating complete mouse embryogenesis. For instance, it is difficult to replicate the later stages of mouse development beyond embryonic day 10.5 (E10.5) in a set-up that allows *in vitro* time-lapse imaging [15, 1].

Hence, in internally developing organisms, it is often necessary to collect and fix the embryos and stage the limbs post-collection. Additionally, biological variability can lead to differences in growth rates even between embryos from the same litter [3]. Accurate staging is thus essential for meaningful comparisons and understanding the temporal dynamics of gene expression during limb development. This requires robust methods for staging and aligning limb buds to ensure that comparisons are made between equivalent developmental stages [11, 17].

Several tools are available for 3D data analysis, among others, Fiji and Napari, which offer powerful general 3D image analysis capabilities [21, 23]. However, these tools are not specialized for the specific challenges associated with analyzing limb development data. Alternatives like Paraview and Imaris provide advanced visualization capabilities but lack customized features to address the unique problems of limb gene expression studiess [2]. Thus, there is a clear need for a computational framework capable of comprehensively analyzing 3D gene expression data in the context of limb development.

In this paper, we present a novel bioinformatics pipeline designed specifically for 3D visualization and analysis of developing limb buds. This pipeline addresses critical challenges in the field, such as the individual staging of limb buds, aligning or morphing to a canonical shape, and general tasks such as data processing and visualization of different volume channels. The different steps of the pipeline are:

- Data Acquisition: Capturing high-resolution 3D images of limb buds, and gene expresison patterns using imaging techniques such as HCR [4], OPT [22] and light-sheet microscopy [13].
- Data cleaning: Removal of noise and artifacts from the raw data to ensure high-quality input for the next steps.
- Limb staging: Independent staging of limb buds providing the morphological stage [17] of the considered limb.
- Transformation: Alignment or morphing of the limb buds to a canonical shape to facilitate consistent comparisons between samples.
- Visualization: Generating detailed 3D rendering for in-depth inspection and interpretation of gene expression patterns.

A schematic representation of these steps is shown in Fig. 1, providing an overview of the process and how each component integrates to facilitate a comprehensive analysis.

**Fig. 1.**
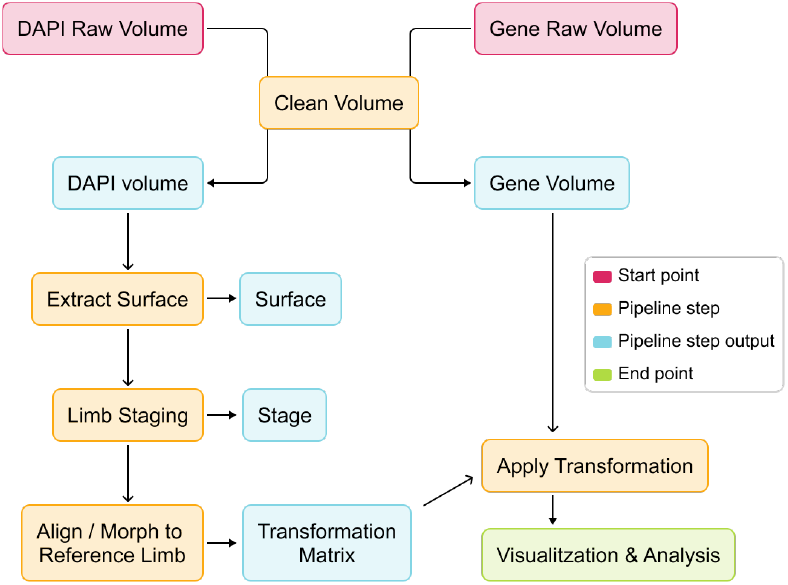
Bioinformatics Pipeline for 3D Limb Bud Analysis. Schematic representation of the pipeline steps: data acquisition (red), data cleaning and transformation (orange), visualization (green), outputs (blue).

This pipeline is designed to be open source and user-friendly, making it accessible to the broader research community. By providing an intuitive interface and comprehensive documentation, we ensure that both experienced bioinformaticians and researchers new to 3D data analysis can easily implement and adapt the pipeline to their specific needs. Using this pipeline, researchers can make more informed decisions, improve the accuracy of their temporal analyses (because of the staging system), and significantly reduce the time required for data processing and visualization. This work introduces significant advances to the current literature by providing a specialized tool tailored to the unique challenges of limb bud development. This novel approach facilitates a better understanding of developmental processes, which can lead to new insights into limb formation and related abnormalities.

## Results

The implementation of our bioinformatics pipeline provides a robust framework for analyzing 3D gene expression data in developing limb buds. This pipeline integrates several crucial steps to facilitate the accurate and efficient analysis of complex volumetric datasets. In this section, we detail each stage of the pipeline. We demonstrate how the pipeline effectively preprocesses raw imaging data, extracts critical anatomical surfaces, and accurately stages and aligns limb buds. In addition, we present advanced visualization tools that allow for an intuitive and detailed exploration of gene expression patterns in 3D. These results underscore the utility of the pipeline in generating reproducible, high-resolution data that ultimately facilitate a deeper understanding of limb development.

### Data pre-preprocessing

The primary requirement of the pipeline is a channel with nuclei labeling - such as DAPI - from which to extract the organ’s outer surface, which is needed for accurate limb staging and alignment to a reference limb. The pipeline assumes that all channels within the same volume are pre-aligned, a condition typically already met by most multi-channel microscopy and mesoscopy techniques.

Before cleaning the volumes, the user is prompted to create a folder, referred to as the *experiment folder*, where all intermediate steps and results will be saved. During this process, the user will also specify whether the limb is *right* or *left, hindlimb* or *forelimb*, as well as the microscope voxel dimensions. Within the experiment folder, a file named pipeline.log will be generated. This file will automatically store all values specified by the user (e.g., clip values used during volume cleaning) and subsequently all automatically assigned values (e.g., the morphological stage).

The user can then begin the volume cleaning process using the command clean-volume. The script first adjusts for the microscope spacing as given in the previous step. The user then selects the bottom and top isovalues to eliminate outliers and reduce noise. Subsequently, the script applies Gaussian smoothing to reduce high-frequency noise, followed by a low-pass filter to remove residual artifacts (Fig. 2 A, B and Movie 1). The refined data are then saved as a vti format file in the experiment folder assigned by the user, ensuring that the volume is ready and easily accessed for further analysis.

**Fig. 2.**
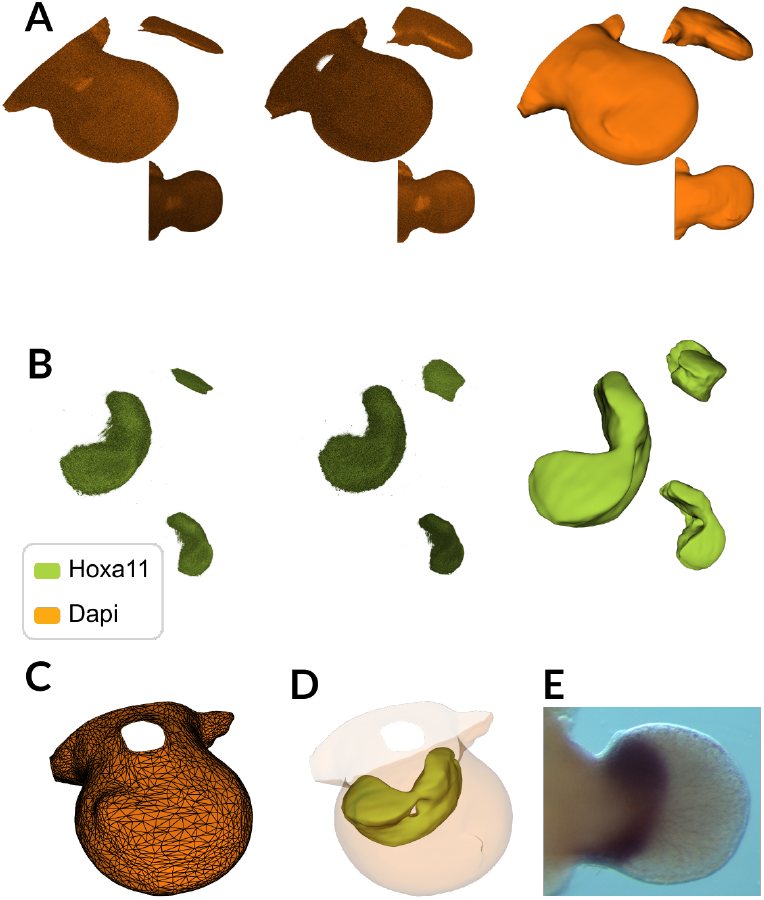
Data Cleaning and Surface Extraction. **A**. Representation of the cleaning steps of a limb bud volume using the DAPI channel. First column shows the row data, second column shows the data after correcting for the voxel spacing, third column shows the volume after the noise reduction. **B**. Same process show in (A) but for the channel of the Hoxa11 gene. **C**. Representation of the extracted surface of the limb. **D**. Visualisation of the limb surface extracted from the from DAPI volumne (light orange) and an isosurface of Hoxa11 gene volume (green). **E** Whole-mount in situ hybridization image showing the expression pattern of the Hoxa11 gene in a developing mouse limb bud.

### Surface Extraction

To optimize staging and alignment, the surface of the organ is extracted as a polygonal mesh, which is computationally more efficient than aligning or morphing raw volume data in the DAPI or equivalent channel. Users can select the isosurface for the outer limb surface either programmatically or through a user interface (UI). This step can be automated, as the pipeline is capable of selecting the most frequent isovalue for extraction, which is then stored in the pipeline.log file. After setting the isovalue, the corresponding surface is extracted. The script subsequently selects the largest connected region, decimates the mesh to reduce its complexity, and optimizes it to minimize noise for downstream analyses (Fig. 2 C, D and Movie 2).

### Staging of the limb

The user can stage - that is, determine the developmental age of a limb - using the command stage. This process involves the manual identification of the Apical Ectodermal Ridge (AER) using the interactive plotter (Movie 3). Once the selection is made, the system sends an API call to the staging server [17]. The server returns the assigned limb morphological stage, along with the accuracy of the staging and a graphic summary of the process.

The morphological stage is saved for use in the subsequent steps of the pipeline (Fig. 3A and Movie 3).

**Fig. 3.**
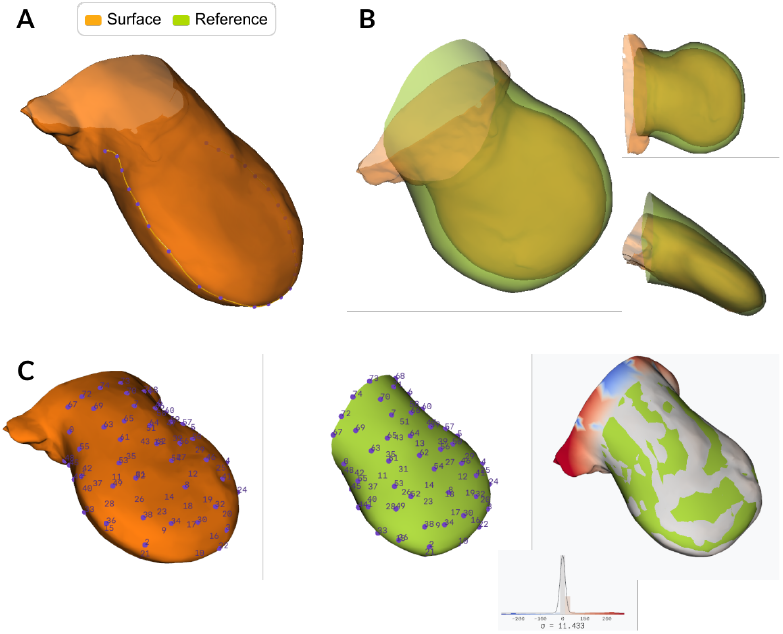
Staging, Aligning and Morphing of the limb. **A**. Visualisation of the interactive plotter for the manually selection of the Apical Ectodermal Ridge (AER). The user must click the limb surface to select the points (purple) that determine the spline of the AER curve (yellow). **B**. Interactive alignment of the limb (orange) to the reference (green) using linear transformation. The user can manually move and rotate the limb to align it with the reference. **C**. Interactive non-linear transformation tool designed to morph a limb to match a reference. Users select pairs of corresponding points (purple) between the limb and the reference. Once a sufficient number of pairs are selected, the tool can automatically generate additional point pairs based on the closest distances. These points collectively define the non-linear transformation used for morphing. Upon completion, the third column displays the distance between the two meshes, providing a visual representation of the alignment accuracy.

### Aligning/Morphing of the limb

With the align command, the user can align the data to a reference limb [7, 8]. This alignment can be performed in one of two possible ways. The first method (command: align) involves a linear transformation that allows the user to rotate and scale the limb interactively while preserving the shape of the surface (Fig. 3 B and Movie 4). The second method (command: align --morph) uses a non-linear transformation, where the user morphs the limb by selecting corresponding points on the surfaces of both limbs. Initially, the user selects a few key points to achieve a rough alignment, after which the script automatically refines the alignment by matching the closest points between the surfaces. This approach ensures precise alignment by closely aligning the shape of the limb with the reference or target limb (Fig. 3 C and Movie 5). Although the alignment process utilizes the surface, the resulting transformation can be applied to the entire volume, ensuring consistency across all data channels. For both methods, the transformation is stored for subsequent use in the aligning of the gene expression pattern volumes.

### Analysis and Visualization

The LimbLab package provides a comprehensive suite of visualization tools designed to accommodate various research objectives, allowing precise exploration and analysis of 3D gene expression data. All of these visualization capabilities are built on top of the Vedo library, a versatile and powerful tool for scientific analysis and 3D visualization [18]. In addition, this package is designed with flexibility in mind, making it easy to add new visualization methods as research needs evolve. All visualitzations are shown in Movie 6.

#### Slices

Displaying slices along the x, y, and z planes allows the researcher to examine cross sections of 3D data at various depths, providing a better view of internal structures and spatial distributions. The detailed 2D slices along these planes help reveal gene expression across different layers of the biological sample. Interactive sliders enable users to dynamically adjust slice positions, making it easier to focus on specific regions of interest. An example of the visualization of the BMP2 gene using this method is shown in Fig. 4A. Additionally, it is possible to visualize the pairing of different genes and extract arbitrary slices of multiple channels. This is shown using the genes BMP2 and *Sox9* in Fig. 4B.

**Fig. 4.**
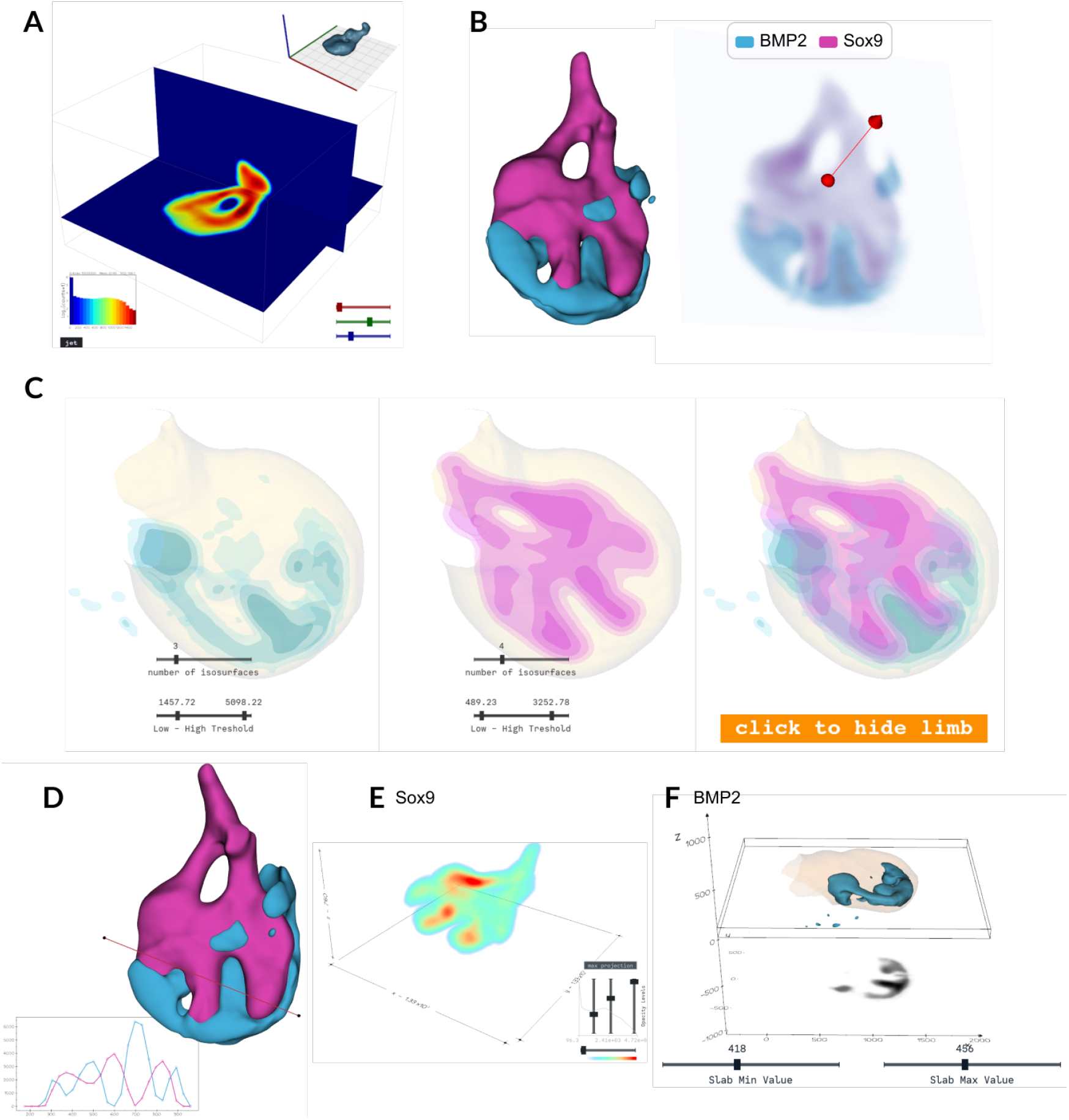
Analysis and Visualization. **A**. Slicing of the 3D expression of the *Sox9* gene (top right) along the x, y, and z planes. The color represent the intensity values of gene expression, which are also depicted in a histogram (bottom left). The user can change colormaps. Sliders allow interactive movement of the x, y, and z planes (bottom right). **B**. Visualization of the genes BMP2 and *Sox9* (left) and an arbitrary slice of these two channels (right). **C**. Isosurface visualization of the volumetric data of a limb (beige) and the expression of the genes BMP2 (blue) and Sox9 (pink). Various sliders enable interactive changes to the number of isosurfaces and the lower and upper threshold values (bottom). A button allows hiding and showing the limb isosurface (bottom right). **D**. Probing two volumes of gene expression, BMP2 (blue) and *Sox9* (pink), with a line, alongside a plot of their respective intensity values. **E**. Raycast visualization of the *Sox9* gene, with a color scheme representing gene expression levels. Sliders adjust opacity, color mapping, and gene expression levels (bottom right). **F**. Representation of a slab (black) extracted from BMP2 volumetric gene expression (blue). Sliders control the slab’s minimum and maximum values (bottom).

#### Isosurface Visualization

Isosurface visualization represents three-dimensional volumetric data by plotting surfaces through regions of constant value. This method effectively visualizes gene expression patterns by allowing users to delineate areas where expression meets specific thresholds. By generating isosurfaces at different values, users can clearly identify and analyze the spatial distribution of gene activity within limbs or organs, facilitating a more intuitive understanding of complex 3D biological data. A key advantage of isosurface visualization is its ability to display multiple gene expression patterns simultaneously in the same 3D space. The overlay of isosurfaces for different genes enables direct observation of spatial relationships, colocalization, and interaction patterns between genes (Fig. 4C).

#### Probe

Probing a gene expression volume with a line and plotting intensity values provides another way of analyzing gene activity in 3D. This involves drawing a line through the volume to examine variations in gene expression, allowing researchers to gain insights into spatial patterns and specific regions of interest. Furthermore, the same line can be applied across different channels, enabling comparative analysis across channels and datasets (Fig. 4D).

#### Raycast Visualitsation

Raycasting is a powerful technique for visualizing 3D volumetric data, such as gene expression patterns, obtained by simulating rays passing through the data to reveal internal structures. LimbLab also includes controls for adjusting the visualization, such as sliders for opacity and color mapping, allowing users to fine-tune how gene expression levels are represented in 3D space. The opacity sliders adjust the transparency of different volume parts, enabling users to focus on areas of interest by making other regions more transparent. Color mapping further enhances visualization by applying various color schemes to different gene expression levels, highlighting specific gene activity thresholds (Fig. 4E).

#### 2D Projection Slab

Extracting a slab from a volumetric dataset simplifies gene expression analysis by converting a 3D volume to a 2D thick slice. This technique combines slices along an axis to create a single image, making it easier to examine gene expression within a specific spatial range. Focusing on a slab helps identify patterns and variations in gene activity, which is particularly useful for studying specific developmental stages or areas with significant changes. Projecting the slab using mean projections highlights different aspects of gene expression: the mean projection shows the average levels, while the maximum projection emphasizes the highest activity. The metadata stored with the slab object, including the range, axis, and operation of the slab, supports a precise and reproducible analysis. (Fig. 4F). To ensure that the slab is perpendicular to the limb, the gene expression volumes and the surface are rotated to correct for variations in the angle of limb protrusion relative to the embryo’s flank [7].

## Conclusion and Discussion

The bioinformatics pipeline presented in this work represents a significant advance in the field of 3D gene expression analysis, particularly in the context of limb development [27]. By integrating a comprehensive suite of tools for data acquisition, volume cleaning, surface extraction, staging, alignment, and visualization, this pipeline addresses the complex challenges inherent in analyzing 3D limb bud data [6, 14, 24, 31]. Its modular design ensures flexibility, allowing users to tailor the workflow to their specific research needs while maintaining compatibility with widely used tools (e.g., Fiji [21] and Vedo [19]).

One of the key strengths of this pipeline lies in its specialization. While general tools like Fiji offer broad applicability, they often lack the convenience and specificity required for niche applications. Our pipeline, on the other hand, is purpose-built to tackle the specific challenges of limb development studies, such as accurate limb staging and alignment, which are critical for meaningful comparative analyses. The ability to seamlessly integrate high-quality visualizations, including isosurface visualization, raycasting, and various projection techniques, further enhances its usefulness, making it a powerful tool for both data exploration and presentation.

A user-friendly interface and robust documentation make the pipeline accessible to researchers with varying levels of expertise, thereby lowering the barrier entry point for complex 3D data analysis. Although the current implementation is optimized for mouse limb buds, the surface extraction and visualization capabilities of the pipeline are applicable to a wide range of biological samples, offering the potential for a wider impact in the field of developmental biology. Moreover, the parameters for visualization parameters, such as color schemes and UI positions are only defined once in the code, hence, it streamlines the creation of concisten publication-ready figures, saving valuable time and effort to researchers.

To ensure that our pipeline evolves in line with the needs of the limb development research community, we actively seek feedback and new requirements from researchers. This engagement will help us tailor future updates and improvements. Upcoming releases will feature new visualizations and additional customization options based on this feedback. Moreover, we plan to explore the integration of tools for analyzing other model species, such as chick and axolotls, which will extend the utility of LimbLab to other fields like evolutionary development and regeneration. Future developments could include the integration of additional staging systems or reference datasets, as well as enhancements to the current visualization techniques to accommodate even more complex biological structures. As the field of developmental biology continues to evolve, tools like this pipeline will be essential in driving forward our understanding of the intricate processes that govern development.

## Materials and Methods

### Materials

#### Embryo Collection

C52BL/6J mouse females were bred and humanely euthanized at different times after gestation to collect embryos at E10.5, E11.0, E11.5 and E12.0. Embryos were dissected and immediately fixed with 4% PFA overnight at 4°C on a shaker. After that, embryos were washed few times with 0.1% PBT and gradually dehydrated with 25%, 50%, 75% and 100% Methanol dilutions in PBT and stored at −20°C. Euthanasia and embryo collection were performed according to European and EMBL guidelines.

### Methods

#### HCRs

In order to perform the HCR, we followed the protocol exactly as described in [16]. In summary, HCR procotol consists of several steps: fixation, photcochemical bleaching, detection, and amplification of HCR™RNA-FISH, clearing, and sample preparation.

#### Sample Preparation

After clearing samples with Fructose-Glycerol solution for a couple of days, samples were embedded in 1% low melting agarose in 10 mM Tris pH 7.4. The agarose gel containing the sample was then cut into octagonal prism shapes of approximately 1 cm^3^ and placed on a 24 well plate with fructose-glycerol clearing solution in order to clear the agarose. This is incubated for a minimum of 24 hours at room temperature protected from light.

#### Data Acquisition

Light sheet imaging was performed using MuViSPIM from Luxendo. Dual side illumination was performed with the illumination objectives Nikon CFI Plan Apo Lambda 4× and collar was adjusted to 1.47, matching the refractive index from Glycerol-Fructose clearing solution. Fluorescence was then captured with immersion 10X Refractive Index adjustment 1.33-1.51 0.5NA glyc WD 5.5 objective. The block of agarose containing the sample was attached to a 3D printed holder and afterwards was immersed in the cuvette containing Fructose-Glycerol clearing solution.

#### Implementation

We implemented the command-line interface (CLI) using Python [12] with Typer [26]. The CLI integrates with our 3D visualization and plotting tools built on Vedo, for the rendering of volumetric data and interactive analysis [18]. We implemented an API for the Staging system [17], and deployed it publicly at [https://limbstaging.embl.es/api]. The API was developed using Python [12] with the FastAPI [25] framework.

## Author contributions statement

LA-E: Conceptualization, Data Curation, Code Implementation, Methodology, Writing – original draft; H.C.-B: Conceptualization, Data Acquisition, Methodology, Writing – original draft; M.M.: Code Implementation, Methodology, Writing – review and editing; A.M.: Code Implementation, Writing – review and editing; JS: Conceptualization, Supervision, Funding Acquisition, Writing – review and editing; G.D.: Conceptualization, Supervision, Project administration, Writing – original draft, review and editing.

## Acknowledgments

The authors thank the Mesoscopic Imaging Facility from EMBL Barcelona for their access to light-sheet microscopy. Specially to Montserrat Coll-Lladó and Jürgen Mayer for the MuViSPIM training and support. We thank the IT department, particullary Juan Carlos Maya, for their support in deploying the online documentation. We also thank Isabel Poetzsch for the bug catching, and the Animal Facility at PRBB for their support in animal welfare, maintenance, and services provided.

This study and the contributions of authors L.A.-E., H.C.- B, M.M., A.M. and J.S. were supported by funding from the European Molecular Biology Laboratory, and the Plan Estatal grant 5972 from MCIU (Ministerio de Ciencia, Innovación y Universidades).

